# Defining the molecular identity and morphology of *glia limitans superficialis* astrocytes in mouse and human

**DOI:** 10.1101/2023.04.06.535893

**Authors:** Philip Hasel, Melissa L Cooper, Anne E Marchildon, Uriel A Rufen-Blanchette, Rachel D Kim, Thong C Ma, Un Jung Kang, Moses V Chao, Shane A Liddelow

## Abstract

Astrocytes are a highly abundant glial cell type that perform critical homeostatic functions in the central nervous system. Like neurons, astrocytes have many discrete heterogenous subtypes. The subtype identity and functions are, at least in part, associated with their anatomical location and can be highly restricted to strategically important anatomical domains. Here, we report that astrocytes forming the *glia limitans superficialis*, the outermost border of brain and spinal cord, are a highly specialized astrocyte subtype and can be identified by a single marker: Myocilin (Myoc). We show that *Myoc*+ astrocytes cover the entire brain and spinal cord surface, exhibit an atypical morphology, and are evolutionarily conserved from rodents to humans. Identification of this highly specialized astrocyte subtype will advance our understanding of CNS homeostasis and potentially be targeted for therapeutic intervention to combat peripheral inflammatory effects on the CNS.

## INTRODUCTION

Astrocytes are an abundant cell type in the central nervous system (CNS) that perform many homeostatic roles that are critical for development and function of other CNS cells like neurons and microglia^1^. Increasingly, astrocytes are appreciated to fall into many discrete subtypes. Their subtype identity is at least in part associated with the anatomical domain they occupy^1–8^, and is likely a consequence of interactions with local cues from adjacent cells. Astrocyte subtypes adjust their transcriptome and functional outputs to the local demand of neuronal circuits and anatomical domains^1,3,8,9^. Recently, we have shown that the astrocyte subtype identity and anatomical position also modulates their heterogeneous responses to pathological insults^2^.

Despite recent efforts using large scale single-cell and single-nucleus RNA sequencing (sc/sn-RNA-seq), specific subtypes of astrocytes that are regionally-restricted remain elusive, aside from transcriptomic subtleties in gross brain regions like the cortex, hippocampus, and striatum^4,5^. In particular, the molecular identity of astrocytes near blood vessels or near the brain surface that have been described histologically is currently unknown^10,11^. Together, these two subtypes make up the *glia limitans* and can be categorized into *glia limitans vascularis* (GLV) and *glia limitans superficialis* (GLS), respectively^12^. Both the GLV and GLS are pre-dicted to have critical roles in defending the brain from peripheral insults and dysfunction, which occurs in many acute and chronic neurodegenerative diseases and would have catastrophic consequences for CNS cells which exist in a tightly-controlled internal milieu^13^. However, the molecular make-up of the GLV and GLS is unknown and it is unclear whether they form two discrete subtypes.

There are contrasting descriptions of the cellular makeup and structural organization of the GLS. Historically, the GLS has been described as astrocytic processes derived from astrocyte cell bodies within deeper layers of the cortex that created subpial expansions closely associated with the brain surface-apposed basal membrane^10,11^. Histological approaches in human and mouse tissue, however, have highlighted that the brain surface itself is in fact occupied by astrocyte cell bodies, whose processes can extend into the parenchyma, in stark contrast to what has been reported historically^14,15^. Despite their prominent location and historical description however, the gene expression profile of these cells remains unknown.

Here, we use a combination of large scale sc/snRNA-seq and genome-wide spatial transcriptomics analysis, whole brain 3-dimensional light sheet microscopy, and *in situ* hybridization to identify the molecular and morphological make-up of GLS astrocytes in both rodent and human brain. We show that the GLS is occupied by specialized astrocytes and confirm recent evidence that their cell bodies reside on the brain surface and have processes that extend into layer I of the cortex. We show that these astrocytes express a unique cassette of genes and can be identified using a single marker: Myocilin *(Myoc). Myoc* is expressed in astrocytes occupying the entire brain and spinal cord surface in a single layer, is induced during early postnatal development, is not expressed in cortical GLV astrocytes, and defines human GLS astrocytes.

## RESULTS

### A transcriptionally continuous domain wraps the brain surface and delineates brain regions

To understand the molecular constitution of the brain surface, we analyzed genome-wide spatial transcriptomics (Visium) of coronal sections from the adult mouse brain (from Castranio et al.^16^). We identified a cluster that defines the entire brain surface as well as demarcates subcortical brain regions (Fig 1a). Beyond tracing the brain surface, this cluster also separates the hippocampus and cortex from the thalamus (Fig 1a). We also find that this cluster selectively expressed a gene, Myocilin (*Myoc*), that reliably tracks the expression of the spatial cluster (Fig 1a). We analyzed brainwide scRNA-seq data (from Ximerakis et al.^17^, Table S1) to show that *Myoc* is selectively expressed in astrocytes, making up a small population of approximately 2% (Fig 1b). By integrating 13 sc/snRNA-seq data sets collected from the murine cortex, hippocampus, spinal cord and striatum (143,966 astrocytes total, Tables S1,S2), we find that *Myoc*+ astrocytes are rare, with an average of 2.4% across all mouse datasets (Fig 1c,d). *Myoc*+ astrocytes are enriched for a range of classical astrocyte genes, such as *Gfap, Clu, Aqp4*, and *Vim* (Fig 1e,f; Tables S3,S4). *Myoc*+ astrocytes also express genes indicating a baseline ‘reactivity’ response in control *Aldh1l1^eG-FP^* mice, including the interferon-regulated transcript *Ifitm3*, as well as reactive marker genes *H2-D1, A2m, B2m*, and *C4b* (Table S3). This combination of reactive astrocyte transcripts is not expressed at baseline in other CNS astrocytes, suggesting a novel function of GLS astrocytes in the healthy brain, potentially regulating how peripheral inflammatory signals enter the parenchyma.

**Figure 1.**
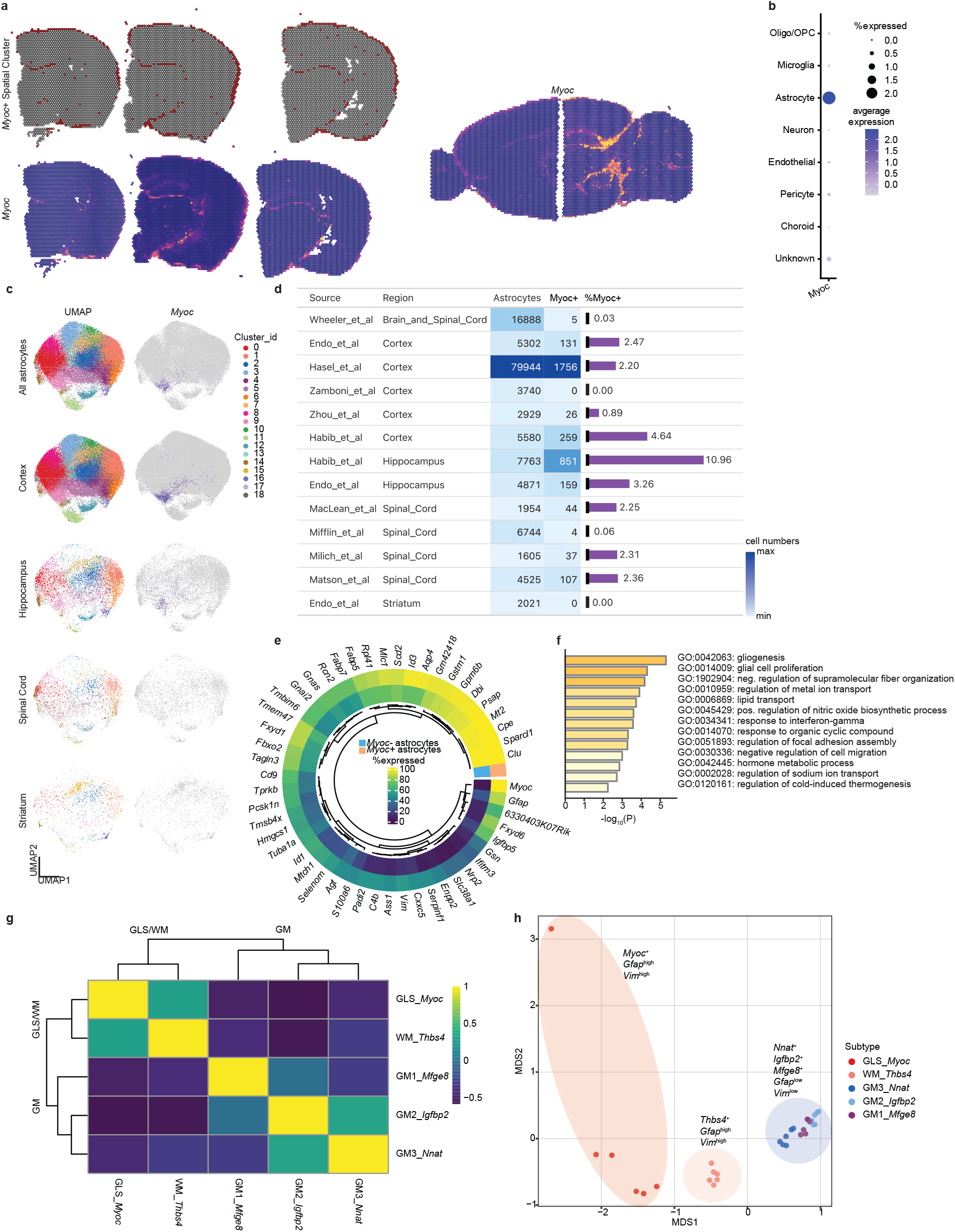
Integrative sc/snRNA-seq and spatial transcriptomics uncovers *Myoc+ glia limitans superficialis* astrocytes. **(a)** Reanalysis of sequencing-based spatial transcriptomics (from Castranio et al.^16^, Table S1) uncovered a continuous transcriptomic tissue wrapping the brain surface and delineating major brain regions defined by the marker Myocilin (*Myoc*). **(b)** sc/snRNAseq analysis from whole mouse brain (from Ximerakis et al.^17^, Table S1) shows that *Myoc* is selectively expressed in astrocytes. **(c,d)** Integration of 13 data sets (from Wheeler et al.^31^, Endo et al.^4^, Hasel et al.^2^, Zamboni et al.^32^, Zhou et al.^33^, Habib et al.^34^, MacLean et al.^35^, Mifflin et al.^36^, Milich et al.^37^ and Matson et al.^38^, see Tables S1,2) across four CNS regions (cortex, hippocampus, spinal cord and striatum) identifies a low number of *Myoc*+ astrocytes (average of 2.4% of all sequenced astrocytes across all regions) in the mouse brain and spinal cord. **(e,f)** Differential gene expression and GO term analysis of scRNA-seq data identifies the transcriptomic identity of *Myoc*+ astrocytes, indicating a white matter-like transcriptome and baseline reactivity. For a full list of genes enriched in *Myoc*+ astrocytes, see Table S3). **(g, h)** Similarity matrix (where are score of 1 describes maximum similarity) and multidimensional scaling plots show that *Myoc*+astrocytes, while transcriptionally distinct, more closely resemble white matter astrocytes (see Fig S1) when compared to grey matter astrocytes (reanalysis from Hasel et al.^2^). GM = grey matter, WM = white matter, GLS =*glia limitans superficialis*. Differential gene expression for both spatial transcriptomics and scRNA-seq was performed using a Wilcoxon rank sum test with adjusted p-values (padj) of <0.05 being considered significant. The sagittal brain sections in a were downloaded from https://www.10xgenomics.com/resources/datasets.

Given the anatomical location of *Myoc*, and its selective expression in astrocytes, we therefore hypothesized that we identified the gene expression profile of, and a selective marker gene for, GLS astrocytes. We also compare *Myoc*+astrocytes with previously reported *Thbs4*+ white matter (WM) astrocytes as well as three subtypes of grey matter (GM) astrocytes (Fig 1g,h; Fig S1). We show that transcriptomically, *Myoc*+ astrocytes align closer to WM astrocyte than GM astrocytes, confirming recent reports that WM astrocytes are transcriptionally more similar to “upper layer” astrocytes when compared to GM astrocytes^2,5^ (Fig 1g,h). Given the apparent absence of Myoc from parenchymal tissue in the spatial transcriptomics data, we hypothesize that these cells specifically represent GLS astrocytes, and that they are transcriptomically distinct from GLV astrocytes.

### The glia limitans superficialis is made up of densely packed cell bodies of Myoc+ astrocytes

In order to confirm that *Myoc* is indeed expressed in GLS astrocytes, we performed RNA *in situ* hybridization (RNA-Scope) of *Myoc* and the astrocyte marker *Slc1a3* (encoding for the Glutamate Aspartate Transporter GLAST). We find that in adult mice (P60), Myoc is indeed expressed in astrocyte cell bodies lining the brain surface, as well as tissues demarcating the cortex and hippocampus from the thalamus, as the spatial transcriptomics data suggested (Fig 2a-d). Myoc+ astrocytes form one layer on each of the internal borders both at the hippocampus-thalamus border, which we argue is the interventricular foramen, as well as the cortex-thalamus border. We speculated that the tissue wrapping the brain will have a similar transcriptomic make up as the surface tissue of the spinal cord. We have detected similarly small numbers of *Myoc*+ astrocytes in sc/snRNA-seq data sets from mouse spinal cord (2.04% of astrocytes are *Myoc*+ in the cortex versus 1.75% in the spinal cord, Fig 1d; Table S2). Our RNA *in situ* hybridization confirms that similar to the brain, the spinal cord surface is occupied by *Myoc*+ astrocytes, which are restricted to the surface and not present around the central canal (Fig 2e,f). On the cortical surface, 99.2% of all astrocytes are *Myoc*+ and 99.8% on the spinal cord surface (Fig 3b). This confirms our hypothesis that *Myoc*+ astrocytes make up the GLS, and that *Myoc* is a reliable marker for the GLS. However, at what point in development the GLS emerges and matures is unclear.

**Figure 2.**
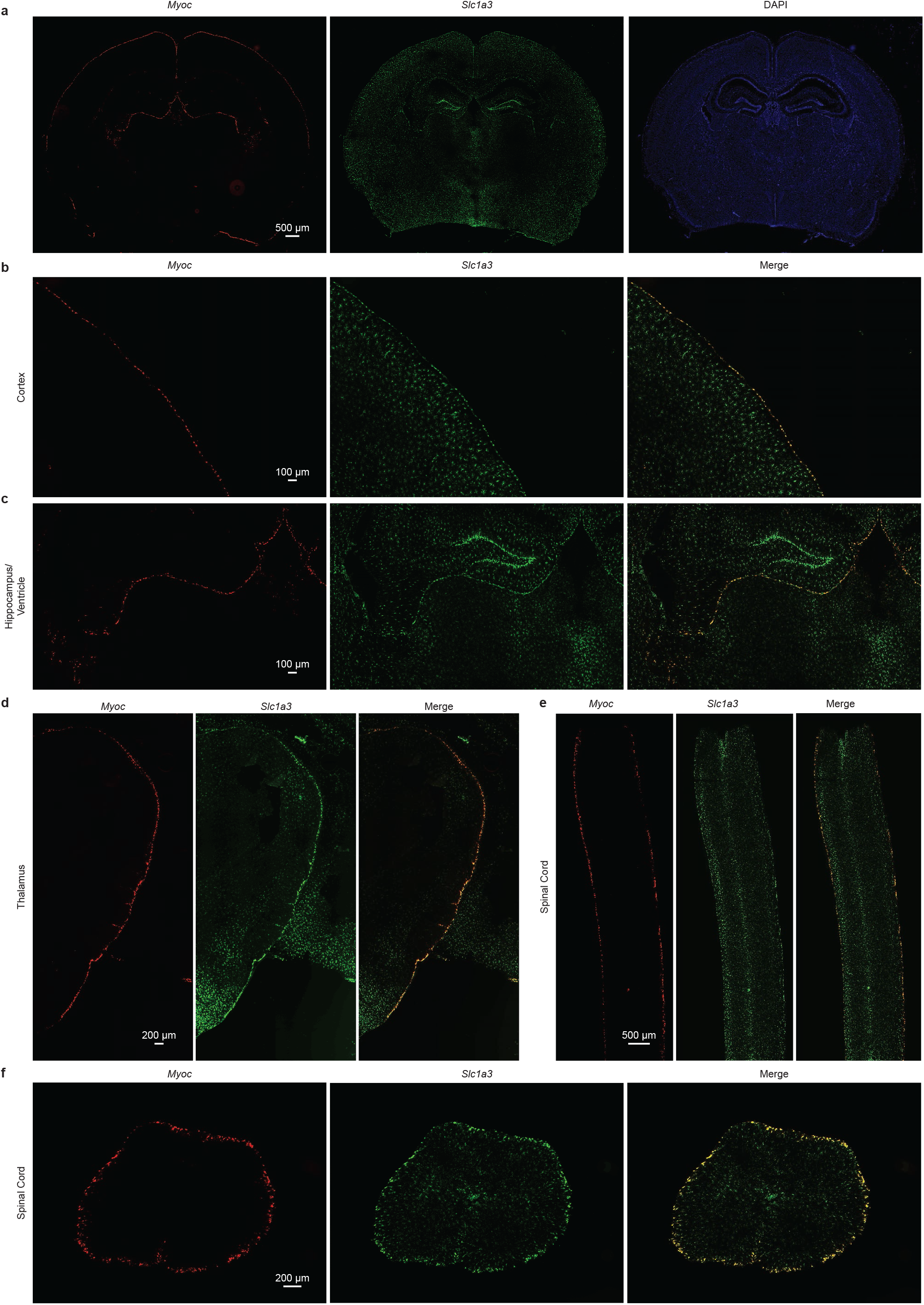
*Myoc*+ astrocytes occupy highly restricted anatomical domains and form the *glia limitans superficialis* in brain and spinal cord. **(a)** Example images of RNA *in situ* hybridization for Myoc and the astrocyte marker *Slcla3* show the highly region-restricted expression pattern of *Myoc*+ astrocytes in the adult mouse brain. **(b)** *Myoc* co-localizes with *Slc1a3*+astrocytes on the cortical brain surface, thereby making up the *glia limitans superficialis* (see quantification in Fig 3). **(c)***Myoc*+ astrocytes can also be observed just ventral to the hippocampus, lining the interventricular foramen and third ventricle. **(d)***Myoc*+ astrocytes also separate the thalamus from the cortex, with one layer on the thalamic and one layer on the cortical side. Unlike in the cortex, in the thalamus, *Myoc*+ astrocytes can also be found wrapping penetrating vessels. **(e, f)** Example images of *Myoc*+ astrocytes in the adult spinal cord showing that *Myoc*+ astrocytes also form the spinal *glia limitans superficialis* by wrapping the entire tissue (see quantification in Fig 3) but are not present in the central canal.

**Figure 3.**
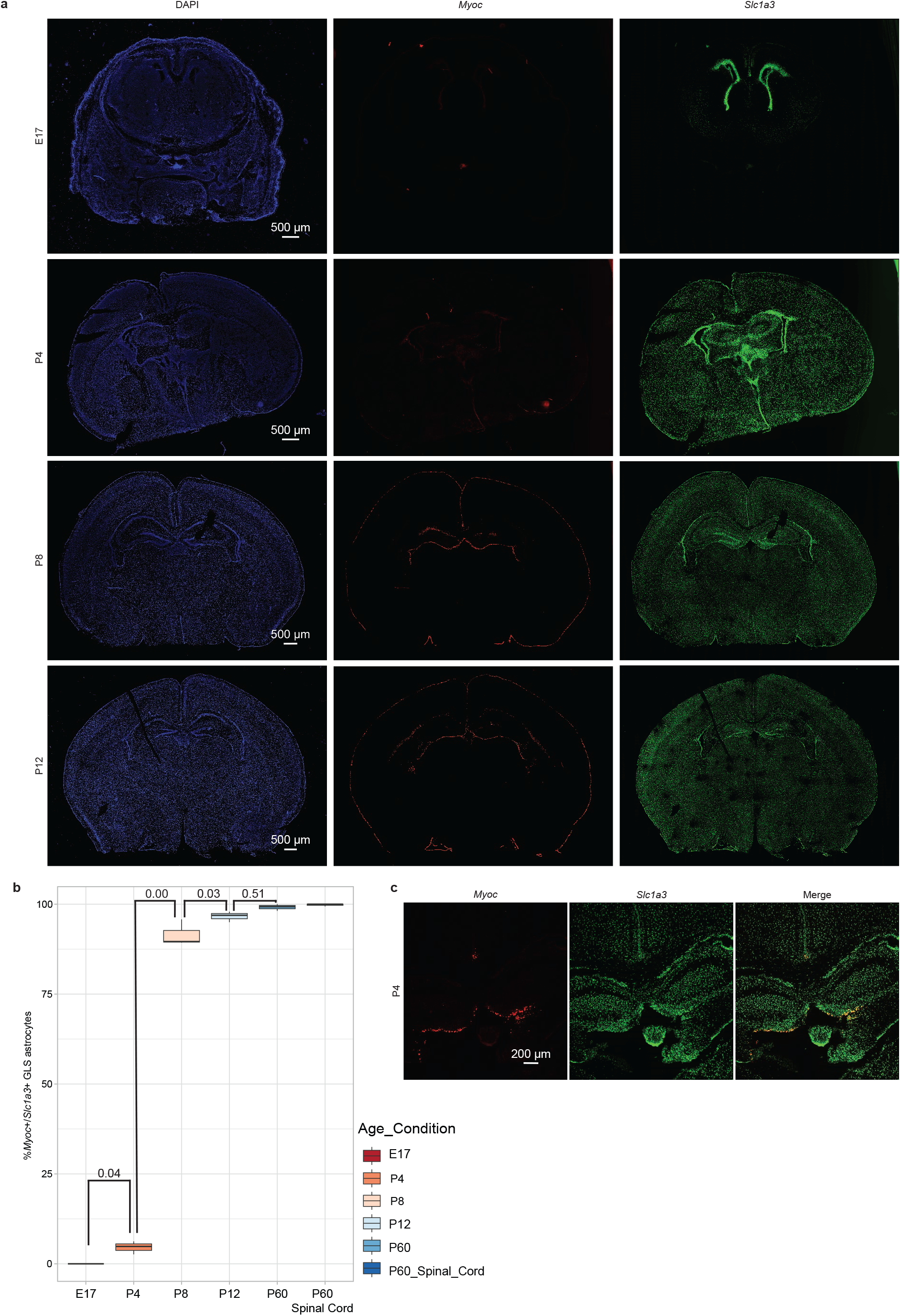
Myoc+ astrocytes of the *glia limitans superficialis* emerge in early postnatal development and are likely instructed locally. **(a,b)** Example *in situ* hybridizations and quantifications for *Myoc* and the astrocyte marker *Slc1a3* throughout late embryonic (E17) and early postnatal (P4, P8 and P12) development as well as quantification of adult (P60) *Myoc+ glia limitans superficialis* (GLS) astrocytes in cortex and spinal cord (n= 3-4 mice per condition). As expected, no *Myoc*+ astrocytes are found embryonically but start to emerge at P4, when 4.6% of all cortical GLS astrocytes are *Myoc*+. This drastically increases to 91.6% at P8 and 96.6% at P12 to ultimately reach 99.2% in adulthood. In the spinal cord, 99.8% of all GLS astrocytes are *Myoc*+. A small subpopulation of white matter astrocytes in the corpus collosum is also *Myoc*+. At no point in development do we observe *Myoc*+ astrocytes in the parenchyma, suggesting that they are instructed locally at the surface. Values in b represent adjusted p-values (see below). **(c)** Example *in situ* hybridization at P4 shows that during early postnatal development, *Myoc*+ astrocytes first emerge just ventral to the hippocampus and that the first *Myoc*+GLS astrocytes mature at the cortical midline. Statistical tests in b were performed using a one-way ANOVA and post-hoc Tukey HSD (Honestly Significant Difference) to calculate padj. A padj of < 0.05 was considered statistically significant.

### GLS astrocytes emerge in early postnatal development and are instructed locally

We next identified temporal dynamics of GLS astrocyte emergence in the developing brain. We collected mouse brain tissue from Embryonic day (E) 17, and postnatal days (P) 4, P8, P12 and P60 and performed RNA *in situ* hybridization. As expected, we found no Myoc+ astrocytes at the embryonic stage (Fig 3a,b). At P4, 4.6% of all cortical *Slc1a3*+ GLS astrocytes are *Myoc*+, initially emerging around the midline (Fig 3a-c). *Myoc*+ astrocytes can also be found emerging just ventral to the hippocampus at the interventricular foramen, an area that similar to the GLS represents a surface domain (Fig 3c). There is a drastic increase of *Myoc*+ GLS astrocytes between P4 and P8, with 91.6% of all cortical GLS astrocytes *Myoc*+ at P8.

By P12, 96.6% of all surface astrocyte are *Myoc*+, nearly reaching levels observed in adulthood (99.2%, Fig 3a,b). We can confirm a similar expression pattern in cortex and spinal cord in adulthood and development in the mouse ISH atlases from the Allen Institute (Fig S2a,b). We see no evidence for transient parenchymal *Myoc*+ cells, indicating that the GLS is instructed locally at the brain surface rather than due to the migration of *Myoc*+ cells to the surface. We speculate that local signals derived from cells located in the meninges could instruct the maturation of the GLS in these early postnatal days. We find a selective ligand-receptor network involving BMP signaling that suggests border-associated macrophages (BAM) residing in subdural meningeal tissue could putatively cause the local maturation of GLS astrocytes (Fig S3). By employing the receptor-ligand analysis tool Cell-Chat^18^ on subdural immune cells (from Van Hove et al.^19^) and GLS as well as sub-GLS astrocytes, we find a putative signaling hub involving BMP2 derived from BAMs activating BMP receptors on GLS astrocytes but not parenchymal astrocytes (Fig S3a-e). Interestingly, BMP signaling has recently been shown to induce Myoc, and other GLS co-expressed genes in primary murine astrocytes, albeit through BMP6 (Fig S3d)^20^. Whether BAM-derived BMP can indeed instruct the maturation of GLS astrocytes remains to be shown. The exact morphology and topology of these *Myoc*+ GLS astrocytes in adulthood, however, is incompletely understood.

### GLS astrocytes cover the cortical surface and have an atypical morphology

In order to profile the morphology and topological organization of GLS astrocytes, we next cleared whole brain hemispheres from adult *Aldh1l1^eGFP^* astrocyte reporter mice with the AdipoClear protocol^21^ and imaged hemispheres intact via light sheet microscopy. Endogenous GFP signals were quenched in the initial bleaching step, then amplified using a chicken anti-GFP antibody (Aves lab, GFP-1020, 1:1000). As our *in situ* hybridization analyses of mouse CNS highlighted that 99.2% of all surface astrocytes are *Myoc*+, we used the *Aldh1l1^eGFP^* astrocyte reporter mouse to study GLS astrocytes by focusing on astrocytes on the surface of the brain (Fig 4a,b,d,e). Light sheet microscopy allowed us to observe the brain surface of an intact brain and compare it to parenchymal tissue (Fig 4b,c). Strikingly, GLS astrocytes possess a morphology entirely unlike parenchymal astrocytes, whose morphology is often described as branched and bushy, with intricate leaflets filling the large volume these cells can occupy^22^ (Fig 4c). In stark contrast, the cell bodies of GLS astrocytes possess a more simple and flat morphology, resembling mono-cultured astrocytes raised in the presence of serum^23^ (Fig 4b,e). Given this atypical flat morphology, it is likely that these cells are often misidentified as subpial expansions from layer I astrocytes, meningeal tissue or mere edge effect. We also confirm that these surface cells extend branched processes into the parenchyma (Fig 4a,d).

**Figure 4.**
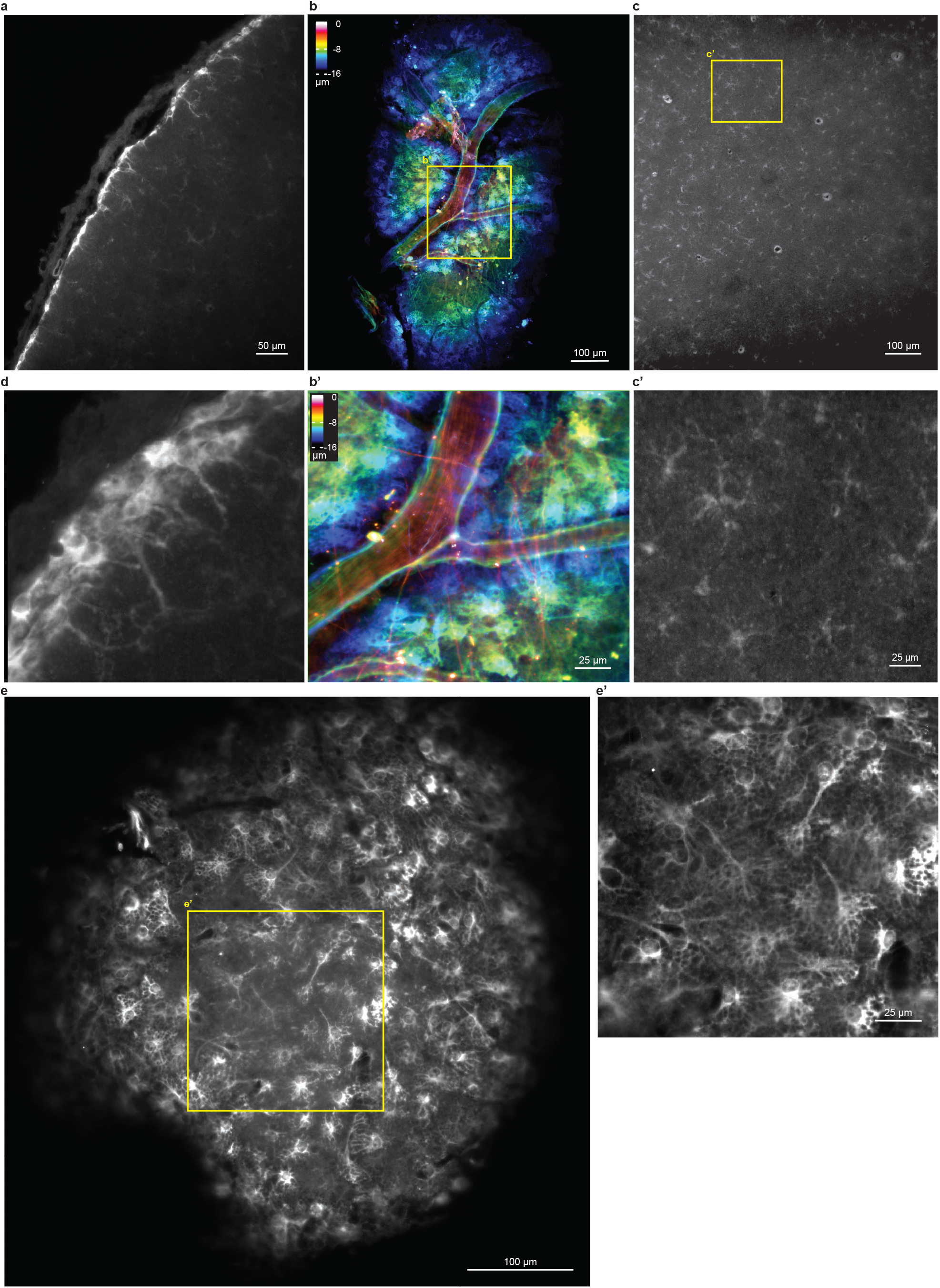
*Glia limitans superficialis* astrocyte cell bodies cover the cortical surface and show an atypical morphology with parenchymal processes. **(a)** AdipoClear-cleared adult *Aldh1l1^eGFP^* brains were imaged using light sheet microscopy. Example images from six animals show astrocyte cell bodies covering the cortical surface, with branched processes extending into the parenchyma. **(b, b’)** Top-down view of the cortical surface shows a superficial blood vessel surrounded by *glia limitans superficialis* (GLS) astrocyte cell bodies. Superficial blood vessels run in GLS ‘grooves’ and are covered in long astrocyte processes that can extend hundreds of micrometers. Color scale represents imaging depth, where 0 μm represents topmost brain structures. **(c, c’)** Parenchymal astrocytes in the same brain show a typical process-bearing, bushy astrocyte morphology. **(d)** Three-dimensional rendering of GLS astrocytes show flat cell bodies occupying the cortical surface and extending branched processes into the brain parenchyma. **(e, e’)** Example images of GLS astrocytes on the cortical surface show densely-packed, process-extending cell bodies.

Like parenchymal astrocytes which cover the entire cortical tissue in a bushy 3-dimensional non-overlapping way, we observed that GLS astrocytes cover the surface of the cortex and arrange themselves around superficial (blood) vessels (Fig 4b,e). At these surface vessel interfaces, GLS astrocytes appear to form ‘grooves’ to accommodate the vessels (Fig 4b). Lastly, we frequently observe long, thin processes that can extend hundreds of micrometers, often superficially crossing superficial vessels, potentially holding them in place (Fig 4b). Having identified the molecular identity and morphology of mouse GLS astrocytes, we next wanted to ask whether Myoc+ astrocytes also make up the GLS in human brains.

### MYOC+ astrocytes occupy the surface of the human brain

Given that Human brains have been reported to also have surface astrocytes with deep penetrating processes^14,24^, we next asked whether *Myoc*+ GLS astrocytes are potentially primitive interlaminar astrocytes and whether *MYOC* is expressed in human GLS astrocytes. To identify *MYOC*+ GLS astrocytes in the human CNS, we integrated recent snRNA-seq data sets from Sadick et al.^25^ and Siletti et al.^26^ (see Tables S1,5) from postmortem human brain tissue. We find that a small population of human astrocytes (on average 0.13% across all data sets and tissues) express *MYOC* (Fig 5a). These *MYOC*+ astrocytes also express similar genes to their mouse equivalent, being high expressers of *GFAP, VIM, CLU*, and *AQP4* as well as the glutamine transporter SLC38A1 (Fig 5b,c; Tables S6,7). Similar to mouse GLS astrocytes, human *MYOC*+ astrocytes express genes traditionally associated with astrocyte reactivity – e.g., compared to all other astrocytes they have increased expression for the inflammatory master regulator *NFKB1*. This suggests that like mouse GLS astrocytes, the CNS surface location presents an environment that maintains an already reactive state under physiological conditions.

**Figure 5.**
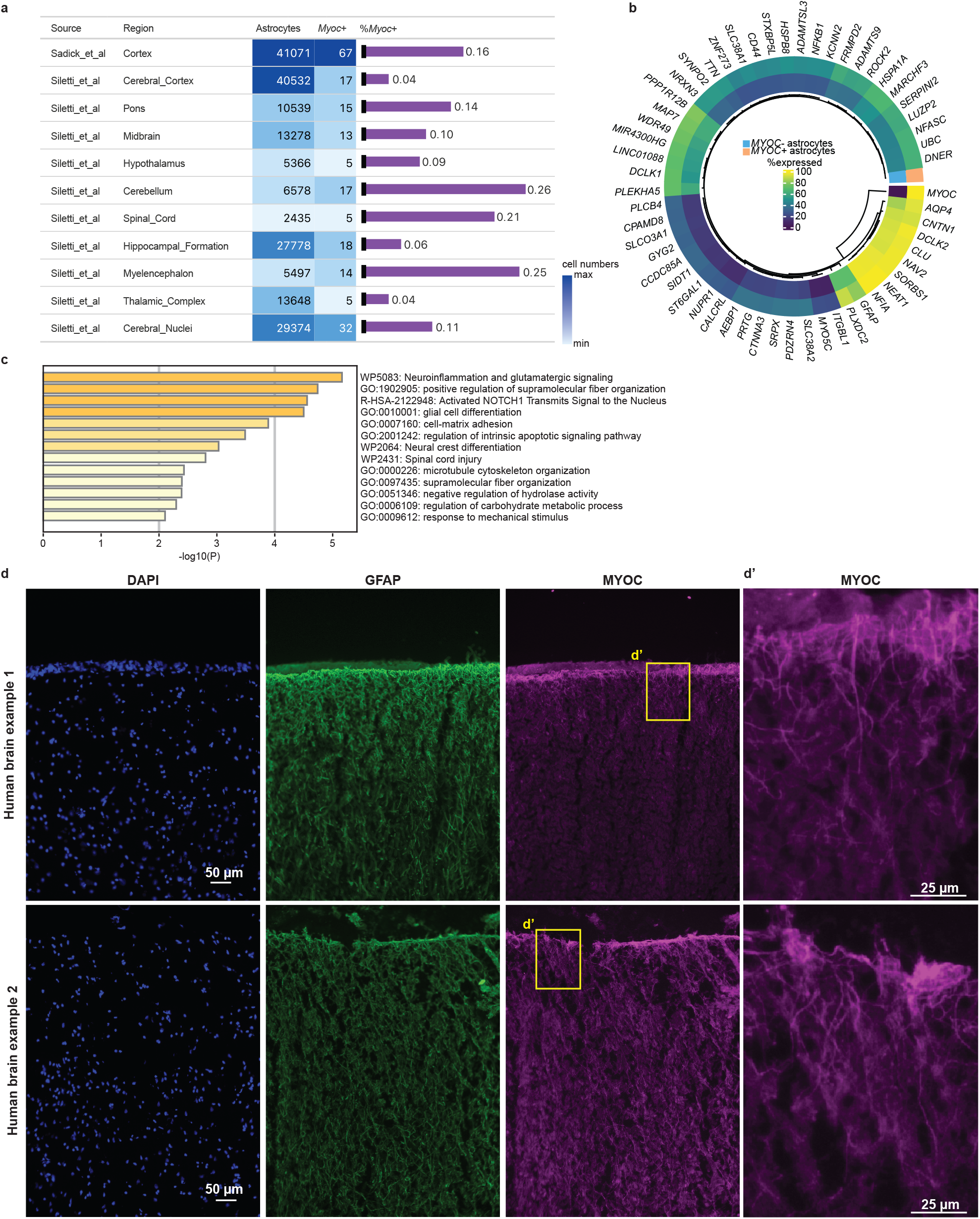
MYOC+ astrocyte cell bodies make up the *glia limitans superficialis* in the human brain and have extensive parenchymal processes. **(a)** Reanalysis of snRNA-seq data from postmortem human brain tissue from Sadick et al.^25^ and Siletti et al.^26^ identifies low numbers of *MYOC*+ astrocytes (0.13% of all astrocytes across all brain regions and data sets, see Tables S1,3). **(b,c)** snRNAseq and GO term analysis identifies the transcriptomic identity of human *MYOC*+ astrocytes. Similar to mouse, human *MYOC*+ astrocytes are enriched in *GFAP, AQP4*, and *CLU* and show indications of baseline reactivity, including enrichment for the transcription factor *NFKB1*. **(d,d’)** Immunohistochemistry identifies GFAP+, MYOC+ astrocyte cell bodies on the brain surface of postmortem human brain tissue with processes that can reach deep into the human prefrontal cortex. Example images from six human brain tissues with identifiable surfaces.

In order to visualize MYOC in human brain tissue, we generated a novel antibody against a small peptide within the olfactomedin domain of MYOC (Fig S4a,b). We used the antibody in postmortem human brain tissue sections and found MYOC+ astrocytes on the surface of the brain creating a mesh-like structure and extending extensive penetrating processes, co-localizing with GFAP (Fig 5d). While we find GFAP+ astrocytes in deeper layers of the cortex, these cells were devoid of MYOC (Fig S5). We argue that MYOC+ astrocytes form the GLS in humans and potentially represent previously described interlaminar astrocytes.

## DISCUSSION

Despite recent efforts in cataloging astrocyte subtypes in health and disease using large scale sc/snRNA-seq, certain histologically-defined subtypes of astrocytes occupying distinct anatomical domains have remained elusive. Highly regionally-restricted and therefore rare subtypes of astrocytes either are not sequenced at all or form too small a population to be identified in standard single cell analysis.

By integrating large numbers of astrocytes derived from a set of mouse and human sc/snRNA-seq data sets of both large and small scope we show that rare but biologically relevant subsets of astrocytes can be computationally identified. By integration of sc/snRNA-seq and genome-wide spatial transcriptomics, we can give a spatial context even to extraordinarily sparse subtypes of cells. Here, we apply these tools to identify the molecular identity of GLS astrocytes in both mouse and human brain, and in mouse spinal cord.

We show that the entire surface of both the brain and spinal cord are occupied by *Myoc*+ astrocytes and thereby make up the *glia limitans superficialis*. We also show that these surface astrocytes have an extraordinary, atypical morphology, with flat cell bodies extending branched processes into the parenchyma, and cell bodies forming grooves to accommodation superficial blood vessels. These cells undergo local maturation at the brain surface during early postnatal development and likely represent interlaminar astrocytes in humans.

What exactly is the role of these GLS astrocytes? Given their high expression of genes encoding astrocyte filamentous proteins, such as Vimentin and Gfap, we speculate that they contribute to the brain’s structural integrity and, using their superficial long, thin processes, hold surface vessels in place. Given their association with superficial blood vessels, they could also be involved in guiding penetrating blood vessels into the brain during development. The deep penetrating processes of GLS astrocytes into the parenchyma likely allow them to communicate with cells in the parenchyma, in mouse particularly cells in Layer I. We speculate these cells are involved in regulating the transduction of inflammatory signals from the periphery and are in a position to quickly communicate the presence of bacterial or viral particles to Layer I neurites, which could affect the behavioral response to the infection. As these cells occupy a highly strategically critical anatomical domain, they will be able to act as‘gatekeepers’ and therefore represent attractive therapeutic targets, though future functional studies to validate such hypotheses will be required.

Given the highly restricted expression of *Myoc* to the GLS in the cortex, we speculate the GLS astrocyte are locally instructed, possibly by leptomeningeal immune cells. We provide evidence for a possible signaling hub from BAMs to GLS astrocytes through BMPs that does not affect parenchymal astrocytes. Cell type-specific manipulation of BMP signaling will uncover how exactly this specialized tissue is instructed during development.

While the function of MYOC is unknown, MYOC is expressed in astrocytes of the optic nerve and optic nerve head, and mutations in *MYOC* can cause glaucoma in humans^27,28^. Whether MYOC plays the same role in the eye and the CNS surface remains to be discovered. How exactly these mutations affect brain and spinal cord function in patients harboring these mutations will shed light on the role of the GLS in disease.

Future research will show exactly how these cells transform in disease. For example, how is the GLS remodeled during stroke or traumatic brain injury where the brain surface is damaged? Is a new GLS formed or does the old GLS re-grow? How is this molecularly orchestrated? How do astrocytes know they are meant to become GLS cells in development and disease? Now that we discovered the molecular identity of GLS astrocytes, we can finally address these fundamental questions. Finally, the presence of MYOC+ GLS astrocytes on the surface of the human brain suggest an important astrocyte subtype that has been evolutionarily conserved. We look forward to ongoing studies of GLS astrocytes in mouse models of disease given their likelihood for translatability.

## METHODS

### Rodent ethics statement

Mice used were either *Aldh1l1^eGFP^* (Tg(Aldh1l1-EGFP) OFC789Gsat/Mmucd, RRID:MMRRC_011015-UCD) or C57BL/6J wild type mice (RRID:IMSR_JAX:000664). Mice had access to both food and water *ad libitum* and housed on a 12-h light/dark cycle. Animal procedures were in accordance National Institute of Health guideline and NYU Langone School of Medicine’s Administrative Panel on Laboratory Animal Care. All animals were housed at 22-25 °C and 50-60% humidity.

### Human postmortem tissue

Deidentified human donor tissue samples from the anterior cingulate cortex were obtained from the New York Brain Bank at Columbia University (n=1 M, 1 F; age: 80-89 years old) and the Neuropathology Brain Bank & Research CoRE at Mount Sinai (n=1 M, 2 F; age: 57-78 years old). Postmortem intervals ranged between 6 and 20 hours. Tissues were fresh-frozen in liquid nitrogen vapor at the time of collection. Aside from one individual with mild hepatic encephalopathy, all subjects had no diagnostic neuropathological abnormalities. All tissue was donated with premortem informed consent and approval from the Institutional Review Board of each institution.

### Meta-analysis of sc/snRNA-seq and spatial transcriptomics data

Spatial transcriptomics and sc/snRNA-seq data sets were downloaded from the corresponding repositories, see Tables S1,2,5. Astrocytes from sc/snRNA-seq data were subset if they expressed all or a critical combination of the following genes: *Gfap, Agt, Aqp4, Apoe, Glul, Slc1a3, Itih3*, and *Aldoc*. Subset astrocytes were then SCTransformed *(Seurat* 4.3.0), merged and variable features from all data sets used for subsequent clustering using *Seurat’s* RunPCA() with 30 PCs, FindNeighbours() and FindClusters() with a resolution of 0.8. Sex was added to vars.to.regress to avoid clustering based on sex-linked genes, such as *Xist*. Data sets were then integrated *(Harmony* 0.1.1) and re-clustered with the same parameters as above. Cells expressing *Myoc* were then subset and FindMarkers() run following PrepSCTFindMarkers() to calculate genes enriched in *Myoc*+ cells.

Spatial Transcriptomics data was SCTransformed, merged and clustered as above using variable features from all brain sections. PrepSCTFindMarkers() was run before FindMarkers() to identify genes enriched in surface spots.

The following packages were used for data analysis and visualization: *Seurat* 4.3.0, *Harmony* 0.1.1, *ComplexHeatmap* (2.14.0), *viridis* (0.6.2), *dplyr* (1.1.0), *ggplot2* (3.4.1), *gt* (0.8.0), *gtExtras* (0.4.5), *svglite* (2.1.1), *circlize* (0.4.15), *ggrepel* (0.9.3), *ggmsa* (1.3.4), *scran* (1.26.2), and *scater*(1.26.1). GO term analysis was run using *metascape*^29^, ligand-receptor analysis was performed using *CellChat*^18^ and pseudobulk-based MDS analysis was run using *muscat*^30^.

### RNA in situ hybridization on mouse brain and spinal cord tissue

Mice were euthanized with CO_2_ at either E17, P4, P8, P12 or P60 and their brains or spinal cords were dissected and placed in cryomolds containing O.C.T (Tissue Tek, 45831EA). The O.C.T. blocks were then placed in dry ice on isopropanol until fully frozen. Samples were stored at −80 °C until sectioning. The tissues were placed in a Lecia cryostat and allowed to equilibrate to −20 °C for an hour prior to sectioning. Tissues were then sectioned on a Leica cryostat at 20 μm onto Superfrost plus microslides (VWR, 48311-703) and stored at −80 °C until use.

RNAScope v2 (ACD) was performed following the manufacturer’s instructions. Tissue sections were taken from the −80 °C freezer and immediately placed in 4% paraformaldehyde in PBS at 4 °C for 15 min. Slides were then rinsed two times with fresh PBS. Sections underwent ethanol dehydration at 50%, 75% and two 100% for 5 min at room temperature. Hydrophobic barriers were drawn around the tissue with an Immedge pen (ACD, 310018). Hydrogen peroxide was added to the sections and left at room temperature for 15 min. The sections were rinsed in water. Protease IV was added to the sections for 30 min at room temperature. The corresponding probes were added: *Myoc* in C1 (ACD, 460981) or *Thbs4* in C1 (ACD, 526821) in combination with *Slc1a3* in C3 (430781-C3). Slides were then incubated for 2 hours at 40 °C in the HybEz oven. Following hybridization, the probes were amplified according to the v2 protocol. Washing was done in between each step with wash buffer. Channel binding was also carried out according to protocol with C1 and C3. Opal fluorophores were added at 1:1000 dilution in TSA buffer. Opal 620 (FP1495001KT) was added to channel 1 for *Myoc* or *Thbs4*, and Opal 690 (FP1497001KT) to channel 3 for *Slc1a3*. Tissue was counterstained with DAPI and mounted with Fluoromount-G (SouthernBiotech, 0100-01) mounting medium and covered with 50 mm microscope cover glass slide (Denville Ultra, M1100-02e. Slides were allowed to dry overnight in the dark at room temperature and subsequently sealed with nail polish. Images were acquired using a Keyence BZ-X710 fluorescence microscope with a 20x objective lens, and images tiled and stitched using Keyence analysis software. To assess the percentage of GLS astrocyte across development, cells were counted with the cell counting feature on Fiji (2.9.0/1.53t). *Slc1a3+/Myoc*+ and *Slc1a3+/Myoc-*cells on the cortical surface were counted for 2-3 brain sections across 3-4 animals.

### Immunohistochemistry on human brain tissue

Human postmortem samples with an identifiable brain surface were embedded in O.C.T., cryosected to 20 μm and mounted on frosted glass slides. Slides were fixed in 4% paraformaldehyde at 4 °C for 20 min, washed in TBS three times and then blocked in 10% normal goat serum (NGS), 0.3% triton in TBS for 1 h at RT. Brain sections were then incubated overnight at 4 °C in rabbit anti-MYOC (1:50, custom) and rat anti-GFAP (Life Technologies, 130300, 1:500) in 5% NGS (MP Biomedicals, 0219135680), 0.3% triton (Sigma-Aldrich, T8787) in tris-buffered saline. Sections were washed three times in TBS and then incubated with goat anti-rabbit Alexa 647 (Thermo Fisher, A-21245, 1:500) and goat anti-rat Alexa 488 (Thermo Fisher, A-11006, 1:500) at RT for 1 h in 5% NGS, 0.3% triton in TBS. Sections were then washed three times in TBS at RT. Lipofuscein autofluorescence was quenched by incubating sections in TrueBlack (Biotium, 23007), sections counterstained with DAPI, covered in mounting media (Fluoromount-G, SouthernBiotech, 0100-01) and cover slipped. Images were acquired on a Keyence BZ-X710 and Keyence software used to tile and stich images.

### AdipoClear and light sheet microscopy of mouse brain

*Aldh1l1^eGFP^* mice were heavily anesthetized with an overdose of pentobarbital (Euthasol: 390 mg Pentobarbital/50 mg Phenytoin/mL at 2 μL/g) and transcardially perfused with PBS and 10mg/L Heparin followed by 4% PFA in PBS. Brains were dissected immediately and post-fixed in 4% PFA in PBS overnight to stabilize glycine residues. Fixed samples were washed in PBS and 0.1% sodium azide 3 times for 1 hour, then stored in PBS/azide until delipidation.

The AdipoClear^21^ protocol was largely followed with a few adjustments specific to brain tissue and this study. To delipidate, fixed samples were washed in 20%, 40%, 60%, 80% methanol in H_2_O/0.1% Triton X-100/0.3 M glycine (B1N buffer, pH 7) for 1 hour each. Samples were fully dehydrated in 100% methanol for three 1 hour washes. Samples were then delipidated in 2:1 dichloromethane (DCM; Sigma-Aldrich): methanol for 1 hour followed by three 1 hour washes in 100% DCM. After delipidation, samples were washed in 100% methanol for 45 minutes twice. To bleach background autofluorescence, samples were incubated in a freshly prepared solution of 80% methanol, 5% H_2_O_2_, and 15% H_2_O. Samples were rehydrated in 60%, 40%, 20% methanol in B1N buffer each for 45 min. Samples were then washed in B1N overnight.

To permeabilize, samples were washed in PBS/0.1% Triton X-100 0.05% Tween 20/2 μg/mL heparin (PTxwH buffer) with an added 45 g/L Glycine and 50 mL/L DMSO. Samples were then washed in PTxwH twice for 1 hour and the brain was separated in to two hemispheres before staining.

Hemispheres were incubated in primary antibody (Aves lab, GFP-1020, 1:1000) diluted in PTxwH for 1 week at 37 °C with rotation. After primary incubation, samples were washed in PTxwH with 8 solution changes over a period of 3 days, then incubated in secondary antibody (Jackson Immuno, 703-606-155, 1:800) diluted in PTxwH for 1 week at 37 °C with rotation. Samples were then washed in PTxwH with 8 solution changes over a period of 3 days.

For tissue clearing, samples were dehydrated in a 20%, 40%, 60%, 80%, 100%, 100%, 100% methanol/H_2_O series for 45 minutes at each step at room temperature. Following dehydration, samples were washed in 2:1 DCM:methanol for 1 hour followed by two 1 hour washes in 100% DCM. Samples were cleared overnight in dibenzyl ether (DBE, Sigma-Aldrich) and stored at room temperature in the dark until imaging.

Whole hemispheres were imaged on a light-sheet microscope (Zeiss Z1) equipped with 5X (used for whole-tissue views with low-magnification) and 20X (used for high-magnification views) objective lenses. Images were collected with two 1920 x 1920 pixels sCOMS cameras. Images were acquired with the Zen Black software (Zeiss). Samples were imaged in a custom imaging chamber filled with DBE and illuminated from both sides by the laser light sheet (light sheet thickness: 4 μm) with a step-size of 1.75 μm.

## Supporting information

Supp_Table_Hasel_et_al_2023

## Acknowledgements

We would like to acknowledge the help of the Microscopy Core at NYU Langone Health with experiments involving light sheet microscopy and image analysis. The computational requirements for this work were supported in part by the NYU Langone High Performance Computing (HPC) Core’s resources and personnel. Funding for this work was provided by NIH/NEI (R01EY033353), the Cure Alzheimer’s Fund, MD Anderson Neurodegeneration Consortium, and The Alzheimer’s Association (SAL); MH119136 and NS107616 (MVC). We also acknowledge the generous support of Paul Slavick and the Parekh Center for Interdisciplinary Neurology (SAL); we would also like to thank the Leon Levy Foundation’s Fellowship in Neuroscience for their support (MLC); we thank Dr Jean Paul Vonsattel of the New York Brain Bank at Columbia University and the Neuropathology Brain Bank & Research CoRE at Mount Sinai for providing human postmortem brain samples.

## SUPPLEMENTARY INFORMATION

### Supplementary Figures

**Fig S1 |** *Thbs4*+ astrocytes reside in the white matter and periventricular domains

**Fig S2 |** Myoc expression according to the Allen Developing Mouse Brain Atlas ISH and Allen Mouse Spinal Cord Atlas ISH

**Fig S3 |** A putative ligand-receptor network exclusive to border-associated macrophages and *glia limitans superficialis* astrocytes

**Fig S4 |** MYOC antibody design and conservation across different species

**Fig S5 |** MYOC is not present in deeper layers of Human tissue

### Supplementary Tables

**Overview |** Supp_Table_Hasel_et_al_2023 includes Tables S1-7.

**Table S1 |** Source of all data sets used throughout the manuscript (alphabetical order)

**Table S2 |** Source and cell numbers of all mouse sc/snRNA-seq data sets used throughout the manuscript

**Table S3 |** Genes enriched in mouse *Myoc*+*glia limitans superficialis* astrocytes

**Table S4 |** GO term analysis of genes enriched in mouse *Myoc+ glia limitans superficialis* astrocytes

**Table S5 |** Source and cell numbers of all human snRNA-seq data sets used throughout the manuscript

**Table S6 |** Genes enriched in human *MYOC+ glia limitans superficialis* astrocytes

**Table S7 |** GO term analysis of genes enriched in human *MYOC+ glia limitans superficialis* astrocytes

**Figure S1.**
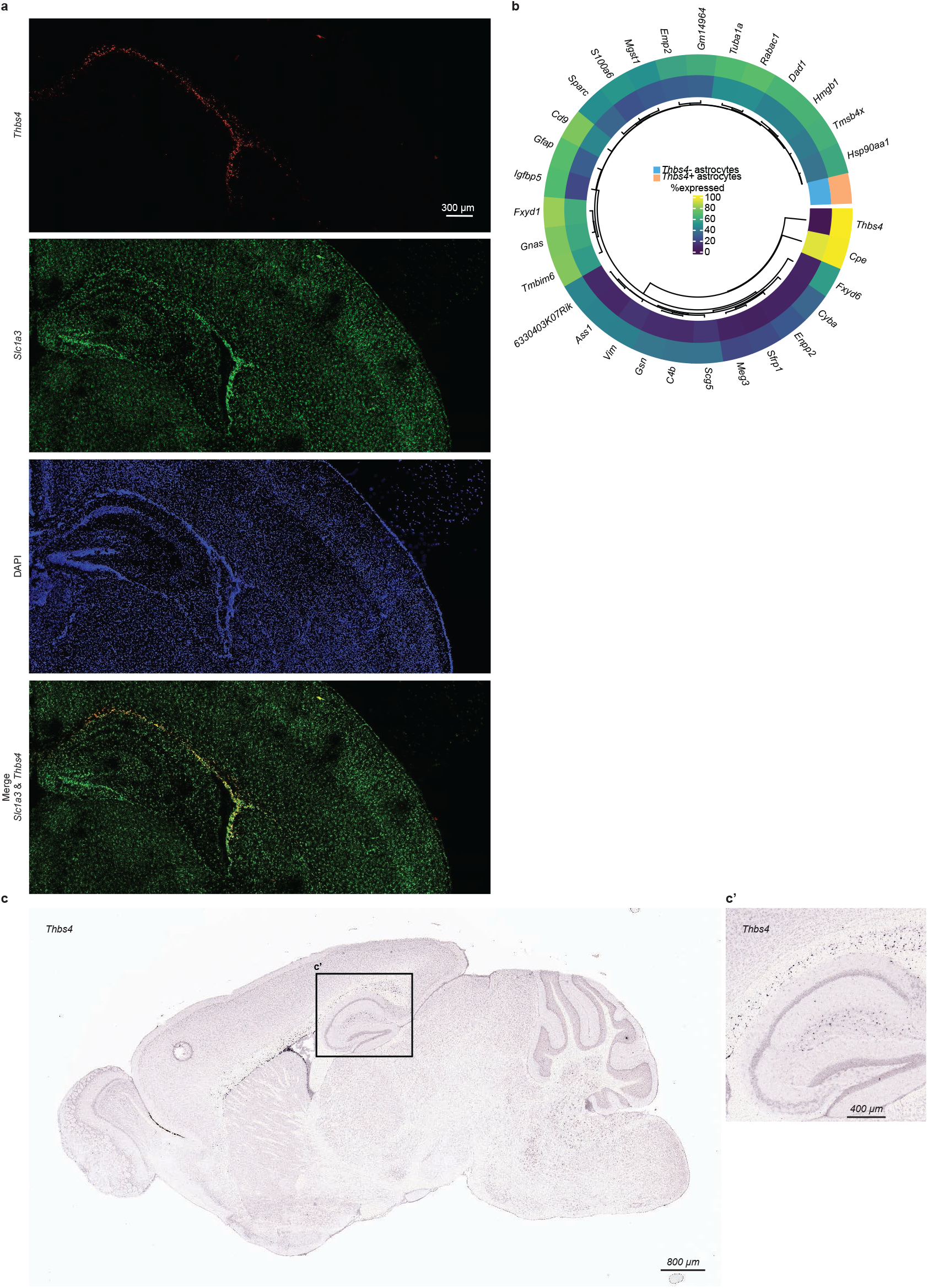
*Thbs4*+ astrocytes reside in the white matter and periventricular domains. **(a)** Example image of *Thbs4+, Slc1a3*+ astrocytes in white matter tracts of a mouse brain. *Thbs4*+ astrocytes track the corpus collosum and surround the third and lateral ventricles. **(b)** Reanalysis of scRNA-seq from Hasel et al.^2^ showing genes enriched in *Thbs4* astrocytes. As shown in Fig 1, *Thbs4*+ astrocytes resemble *Myoc+ glia limitans superficialis* astrocytes, co-expressing *Gfap, Vim*, and both FXYD-domain containing genes *Fxyd1* and *Fxyd6*. **(c,c’)***Thbs4* expression from the Allen Mouse Brain Atlas ISH39 confirms presence of *Thbs4*+ astrocytes in white matter tracts and periventricular domains. From the Allen Institute for Brain Science (2004). Allen Mouse Brain Atlas ISH (https://mouse.brain-map.org/gene/show/21587).

**Figure S2.**
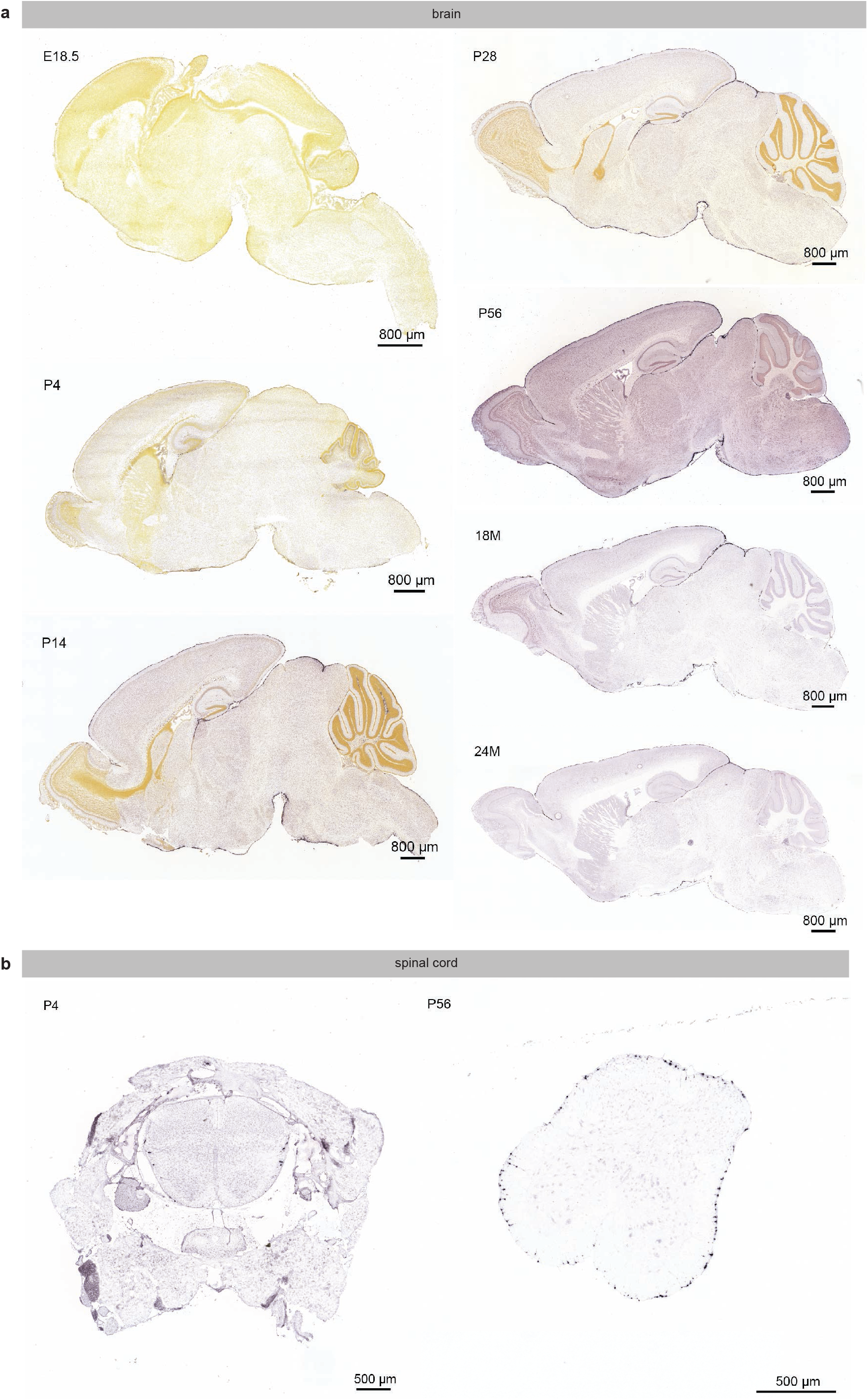
*Myoc* expression according to the Allen Developing Mouse Brain Atlas ISH and Allen Mouse Spinal Cord Atlas ISH. **(a)** *Myoc* expression from the Allen Developing Mouse Brain Atlas ISH^40^ confirms presence of *Myoc*+ cells on the mouse brain surface in development and adulthood. From the Allen Institute for Brain Science (2008). Allen Developing Mouse Brain Atlas ISH (https://developingmouse.brain-map.org./gene/show/17693). **(b)** *Myoc* expression from the Allen Mouse Spinal Cord Atlas ISH^41^ also confirms presence of *Myoc*+ cells on the spinal cord surface. From the Allen Institute for Brain Science (2008). Allen Mouse Spinal Cord Atlas ISH (P56 https://mouse.brain-map.org/experiment/siv?id=100003850&imageId=100061440 and P4 https://mousespinal.brain-map.org/imageseries/show.htm-l?id=100003727).

**Figure S3.**
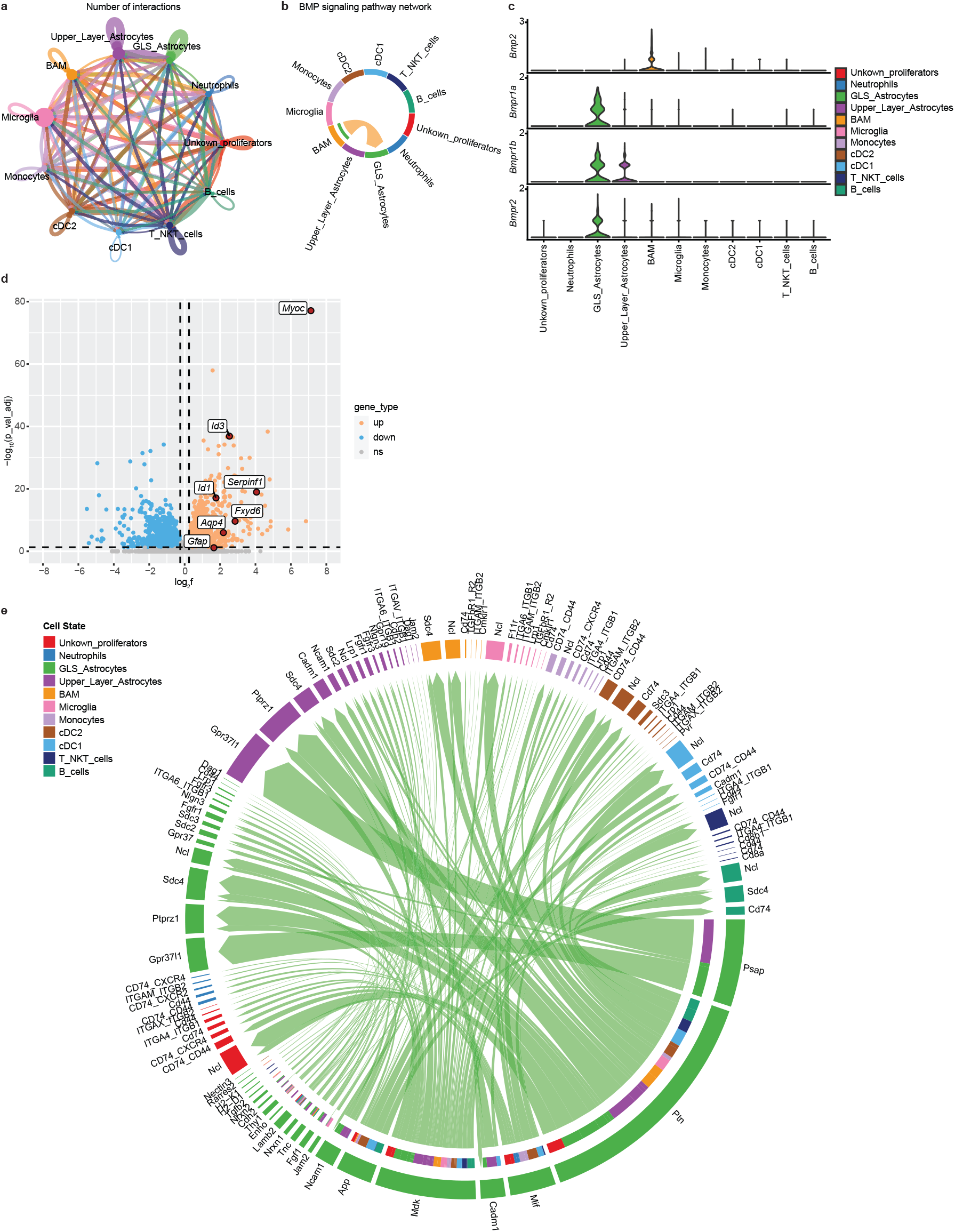
A putative ligand-receptor network exclusive to border-associated macrophages and *glia limitans superficialis* astrocytes. **(a)** Receptor-ligand interactome using CellChat18 on subdural immune cells (from Van Hove et al.^19^, see Table S1) and cortical as well as *glia limitans superficialis* (GLS) astrocytes (from Hasel et al.^2^). **(b,c)** CellChat identifies a putative BMP signa ling network that is exclusive to border-associate macrophages (BAMs) and GLS astrocytes and involves BMP2 release from BAMs activating BMP receptors on GLS astrocytes. **(d)** Reanalysis of Caldwell et al.^20^ shows that BMP signaling, through BMP6, is sufficient to induce *Myoc* and additional genes co-expressed in GLS astrocytes, including FXYD-domain containing gene Fxyd6 as well as Id1 and Id3 in primary, immunopanned murine astrocytes. **(e)** Outgoing signaling from GLS astrocytes to subdural immune cells and cortical astrocytes. GLS =*glia limitans superficialis*, BAM = border-associated macrophages, cDC = conventional dendritic cells, T/NKT cells =T-cells/Natural killer cells.

**Figure S4.**
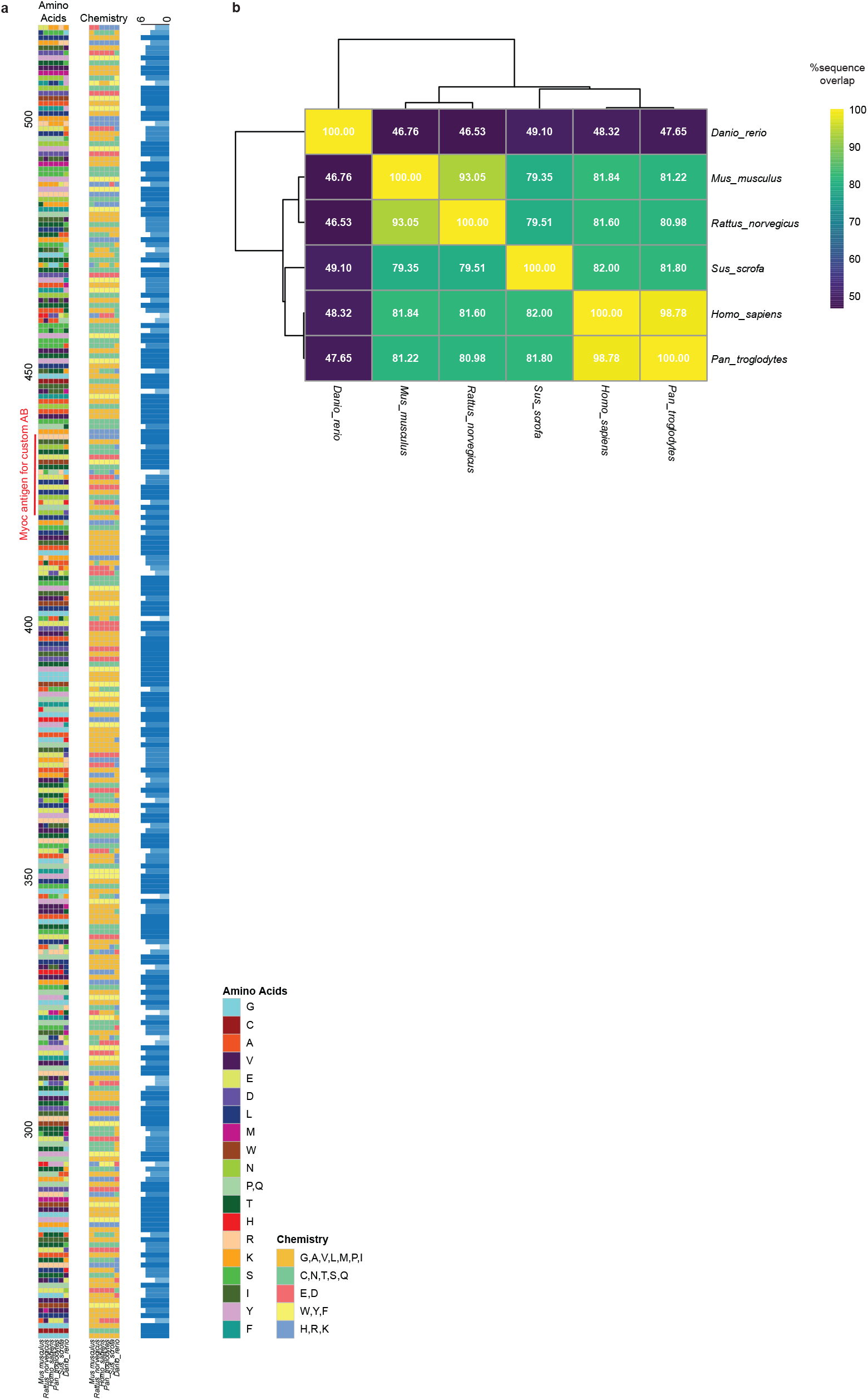
MYOC antibody design and conservation across different species. **(a)** Amino acid sequence containing the olfactomedin domain of MYOC. Highlighted is the peptide sequence used to produce the custom antibody in rabbits. **(b)** Heatmap showing the conservation of the MYOC amino acid sequence across different species.

**Figure S4.**
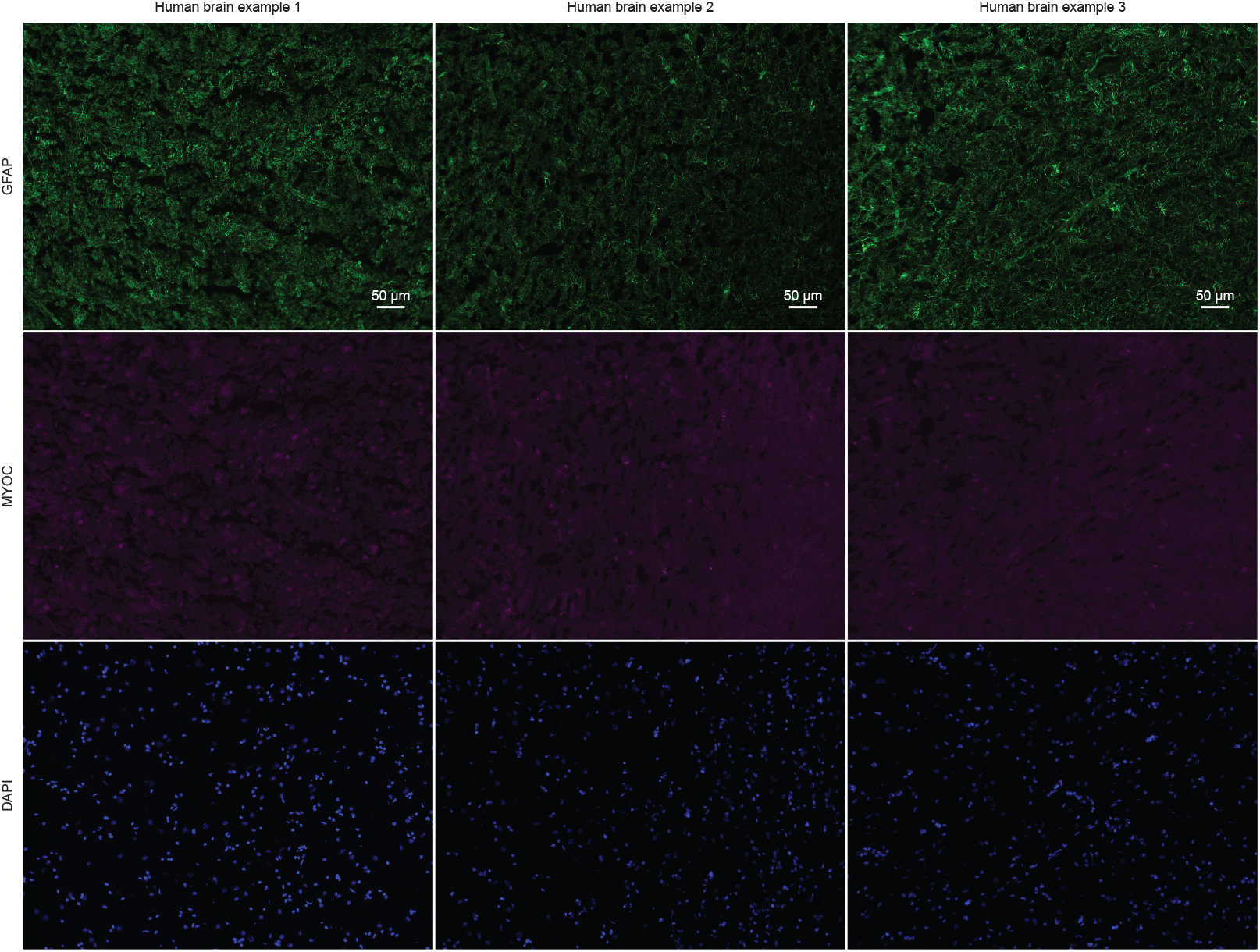
MYOC is not present in deeper layers of Human tissue. Example images of human brain sections used in Fig 5 showing no MYOC immunoreactivity in GFAP+ astrocytes in deeper layers of the human brain.

